# A Dual Reward-Place Association Task to Study the Preferential Retention of Relevant Memories in Rats

**DOI:** 10.1101/2020.01.27.921593

**Authors:** Frédéric Michon, Jyh-Jang Sun, Chae Young Kim, Fabian Kloosterman

## Abstract

Memories of past events and common knowledge are critical to flexibly adjust one’s future behavior based on prior experiences. The formation and the transformation of these memories into a long-lasting form are supported by a dialog between the coordinated activity of population of neurons in the cortex and the hippocampus. Not all experiences are remembered equally well nor for equally long. It has been demonstrated experimentally in humans that memory strength positively depends on the behavioral relevance of the associated experience. Behavioral paradigms testing the selective retention of memory in rodents would enable to further investigate the neuronal mechanisms at play. We developed a novel paradigm to follow the repeated acquisition and retrieval of two contextually distinct, yet concurrently occurring, food-place associations in rats. We demonstrated the use of this paradigm by varying the amount of reward associated with the two locations. After delays of 2h or 20h, rats showed better memory performance for experiences associated with larger amount of reward. This effect depends on the level of spatial integration required to retrieve the associated location. Thus, this paradigm is suited to study the preferential retention of relevant experiences in rats.

## Introduction

Memory is the ability of the brain to encode and store information for later use. The ability to remember past events and facts, is critically dependent on the medial temporal lobe and its connections to the cortex (Squire, Stark, and Clark 2004). Following initial formation (encoding), a memory trace undergoes active post-processing that stabilizes the trace and integrates it into the brain’s existing knowledge base (consolidation). Both encoding and consolidation are supported by the coordinated activity of neuronal ensembles in the hippocampus and cortical areas (Battaglia et al. 2011). Memory consolidation predominantly occurs during sleep. It engages a bidirectional cortico-hippocampal dialogue characterized by the occurrence of cortical slow wave oscillations, spindles and hippocampal sharp wave ripples (SWRs) (Todorova and Zugaro 2018).

However, not all experiences are remembered equally well or for equally long. A growing body of literature in humans has shown that behaviorally relevant aspects of experience, such as emotional content or expected outcome during learning (Payne et al. 2008; Igloi et al. 2015; Wamsley et al. 2016; Studte, Bridger, and Mecklinger 2017), enhance the retention for the associated memory (Stickgold and Walker 2013). Relevant material is preferentially remembered even in comparison to neutral material occurring concomitantly or close in time. Moreover, the enhanced retention of such experiences correlates with the increased hippocampal activity during learning, as well as post learning (Rauchs et al. 2011; Gruber et al. 2016) and with the increased occurrence of slow wave sleep and spindles in the cortex (Stickgold and Walker 2013; Igloi et al. 2015; Gruber et al. 2016; Studte, Bridger, and Mecklinger 2017). Thus, the enhanced retention of relevant experiences is an active selective process relying on the modulation of the neuronal activity supporting both encoding and consolidation.

Rodents are commonly used as animal models to study cognitive functions and in particular the neurobiology of learning and memory. They present a combination of several advantages. First, rodents require relatively low resources to maintain and can be trained in various behavioral assays (Tolman 1948; Hodges 1996; Rosenfeld and Ferguson 2014; Wood et al. 2018). Second, they share anatomical and functional similarities with human, especially for key brain regions supporting learning and memory such as the medial temporal lobe (Eichenbaum, Yonelinas, and Ranganath 2007). Finally, well established tools and techniques exist to monitor neuronal activity, via electrophysiological recordings or optical imaging (Kloosterman et al. 2009; Buzsáki et al. 2015; Ziv and Ghosh 2015; Weisenburger and Vaziri 2018), and to manipulate activity in order to determine causal links with the animals’ behavior (Girardeau et al. 2009; Buzsáki et al. 2015; Latchoumane et al. 2017). Aspects of experience, such as rewarded outcomes, have also been shown to affect memory retention in rodents (Salvetti, Morris, and Wang 2014). However, to our knowledge, no study has reported the selective retention of memory for experiences occurring concomitantly.

We developed a novel paradigm to study the selective retention of memories in rodents in which the retention of two, concomitantly acquired, food-place associations is assessed every day. This behavioral paradigm was successfully used to confirm the improved memory retention of experiences associated with larger reward size, and further, to demonstrate the causal role of post-learning hippocampal replay in the reward-related enhancement of memory consolidation (Michon et al. 2019).

## Materials and Equipment

### Animals

A total of 21 male Long Evans rats were food deprived to 85-90% of their free-feeding weight. Of these, 13 animals received an implant for electrical recording (as part of another study), and 10 rats did not undergo surgical procedures and were only tested behaviorally. All experiments were carried out in accordance with protocols approved by KU Leuven animal ethics committee (P119/2015) in accordance with the European Council Directive, 2010/63/EU.

### Apparatus

The behavioral testing apparatus was located in one of two 4×4 m rooms with black walls and distinctive distal cues on each of the walls. The apparatus was elevated 40 cm of the ground and consisted of a home platform that gave access to the left and right side of the room via a short 30 cm track (figure 1a,b). The left and right environments were separated by 120 cm high dividers. In each environment, a choice platform gave access to a maximum of 6 radially emanating 90 cm long arms separated by 30°. Access from the home platform to the two environments was controlled by a door that was manually positioned by the experimenter. To prevent the animals from using olfactory cues to navigate the maze, food dispensers reward that could be smelled but was otherwise inaccessible to the animals were positioned at the end of every arm. Moreover, the maze floor was covered with rubber sheet that were cleaned with water and pseudo randomly swapped throughout the training sessions.

**Figure 1:**
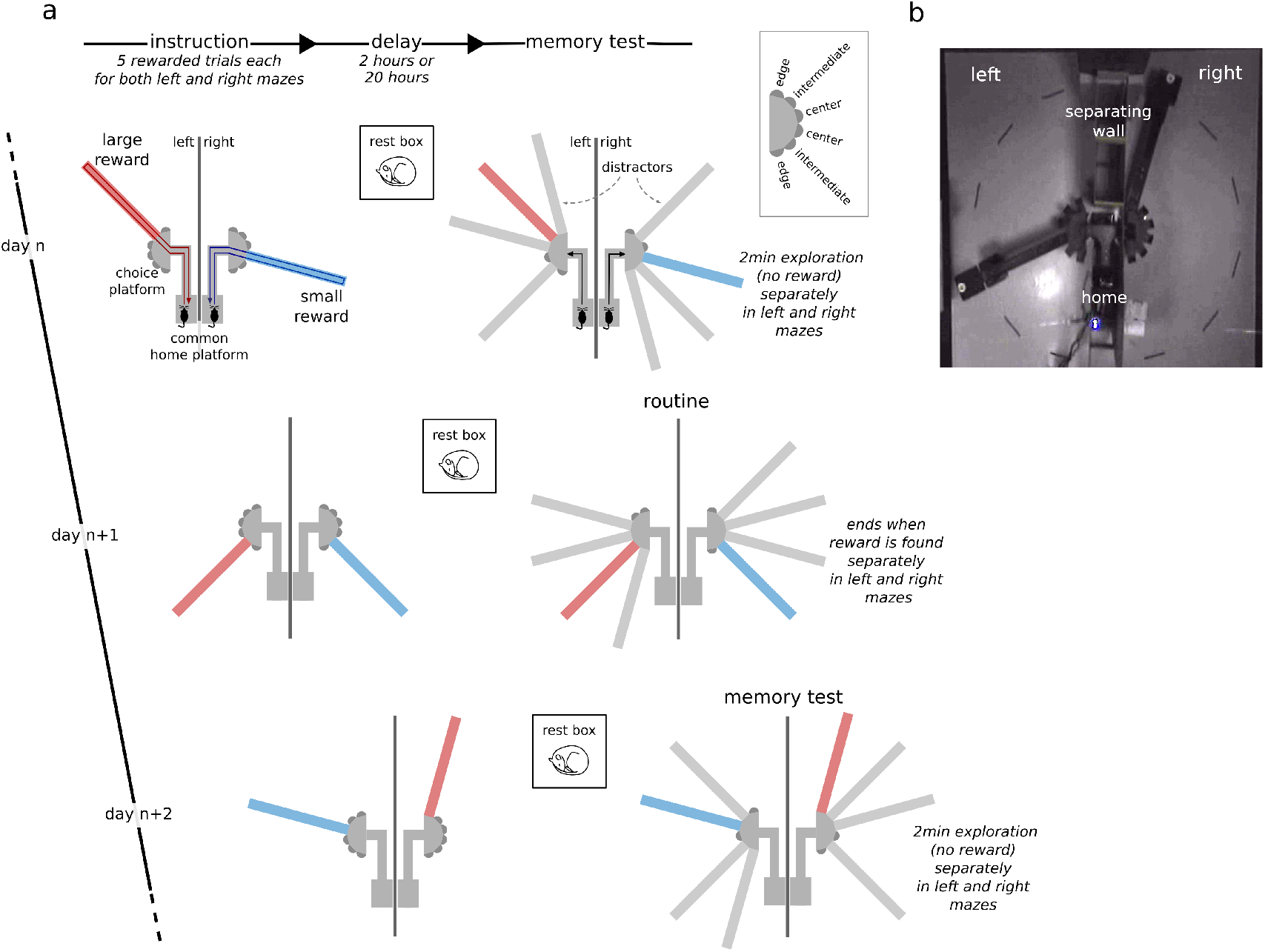
Dual reward-place association task. (a) The behavioral task is composed of three phases: instruction, delay and memory probe test. During instruction, rats learn to associate a small reward (blue) or a large reward (red) with a specific target arm in the left and right environment. During the memory test after the delay, the preference for the target arm in presence of three distractor arms is assessed as a measure of memory. Inset: labels for target arms based on their location relative to the separating wall. The first trial after the delay phase was either a reinstatement trial or a memory probe trial, and this was alternated from session to session. Across sessions, the location of the rewarded target arms, the configuration of the distractors and the small/large reward assignment to the left/right environment were varied pseudo-randomly. (b) Video frame from an overhead camera that shows the apparatus during an example session.

## Methods

### Behavioral task

The goal of the task was for the rat to associate one of the 6 arms in each environment with a reward. In each daily session, one environment was associated with a large reward (9 pellets) and the other with a small reward (1 pellet). After an acquisition phase, during which the animal could explore the rewarded target arms across 5 instruction trials per environment, and after a subsequent 2h or 20h delay, the rats were tested for their memory of the reward-place association in the presence of three distractor arms (figure 1a). Across sessions, the location of the target and distractor arms and the assignment of large/small reward size were varied pseudo-randomly (figure 1a).

### Behavioral procedure

Prior to experimental sessions, the animals were gently handled by the experimenter at least 5 minutes per day for 7 days and pre-trained to run back and forth on an elevated linear track (40cm high and 90cm long) to obtain food rewards (3 pellets) until the animals executed at least 20 laps within 10 minutes for three sessions in a row. During this phase the animals were also habituated to being constrained every time they reached one end of the maze by a door manually controlled by the experimenter.

Next, the rats were familiarized with the experimental procedure and the maze environment of the dual reward-place association task. During the instruction phase, only the two target arms were physically present in the two environments. The instruction phase consisted of 5 blocks of alternating trials to the right and left environment. Each trial began with the animal constrained to the home platform. It was then given access to only one of the two environments. The following trials started after the rat had consumed the reward at the end of the target arm and returned to the home platform. The presentation order of the environments within the trial blocks was constant within a session and randomized across sessions. After the instruction phase and before the test phase, the rat was removed from the maze apparatus for a short (at most 15 minutes) delay and kept in his home cage. After the delay, rats were subjected to three reinstatement trials separately for the two environments in the presence of the distractor arms. In each reinstatement trial, rats were rewarded for visiting the target arm with 3 pellets. The aim of the reinstatement trial was for the animal to learn to seek for a reward at the end of the target arm and to ignore the distractor arms. Each reinstatement trial lasted until the animal consumed the reward at the end of the target arm.

The pretraining phase ended when the rats first visited the target arm during the test in both environments for three days in a row. During the experimental phase, the rules of the dual reward-place association task and topography of the maze were kept the same, but different reward sizes (1 and 9 pellets) and longer delays (2h or 20h) were introduced. During the delay phase, the rat was either returned to its home cage or placed in a 40×40 cm rest box with 60 cm high walls that was located inside the behavioral room. In one out of every two sessions, the first reinstatement trial was replaced by a two-minute-long unrewarded memory probe trial separately for the large and small reward environments. After pauses in training (e.g. during weekends), the subsequent experimental session was preceded by a pre-training session. This procedure was followed to make sure that the rats retained their motivation to search for reward at the target arm in the memory probe trials.

### Data analysis

Data analysis was performed using Python (Millman and Aivazis 2011) extended with custom toolboxes.

#### Behavior

In the instruction trial, the average running speed to and from the reward platform was quantified only for implanted animals, based on video tracking of the headmounted LEDs. Average speed was computed over the journey that started when the animal left the home platform and ended when the animal reached the reward platform at the end of a target arm (and vice-versa for the homebound journey).

In the memory probe trial, the number and pattern of visits to the target and distractor arms were quantified as measures of performance in the reward-place association task. A visit to an arm was only counted if the animal reached the reward platform at the end of an arm. We defined the following quantities and (conditional) probabilities to characterize the reward-seeking behavior in the probe trial:

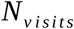

the total number of arm visits in the 2-minute memory probe trial.

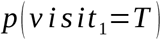

the across session mean probability that the first visit is on target.

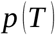

the across session mean probability that a visit is made to the target, computed by averaging the equivalent per session *p*(*T*). This probability is further split in the conditional probabilities *p*(*T* ∨ *D*) and *p*(*T* ∨ *T*) that measure the mean probability of visiting the target arm given that the immediately preceding arm visit was also on target (*p*(*T* ∨ *T*)) or was to a distractor (*p*(*T* ∨ *D*)).

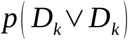

the across session mean probability that a repeat visit is made to any one of the three distractor arms, computed by averaging the equivalent per session *p*(*D_k_* ∨ *D_k_*).

#### Statistics

To test a difference in means between two paired samples, we used a Wilcoxon signed rank test. To test for a difference in two sample proportions, we used either the McNemar test (for paired samples) or the two-proportion z-test.

To analyze the dependence of behavioral metrics on predictor variables, we fitted Bayesian generalized linear models (GLMs) using the PyMC3 package for Python (Salvatier, Wiecki, and Fonnesbeck 2016). We applied a Poisson regression model (with log link function) for the number of arm visits, a logistic regression model (with logit link function) for *p*(*visit*_1_ = *T*) and an ordinary linear regression model for *p*(*T*).

Model fitting and inference was performed using Markov chain Monte Carlo (MCMC) sampling methods in PyMC3 (specifically, the No-U-Turn Sampler). Broad normal distributions were used as priors on the parameters.

## Results

### Fast learning of reward-place associations

In the instruction phase of the task, we first asked whether the behavior of the rats differed between large and small reward instruction trials, as evidence of fast acquisition of the association between reward magnitude and targets in left/right environment. Indeed, the average running speed towards the reward platform was significantly higher in instruction trials for the large reward amount as compared to the small reward amount (figure 2a, left; mean [99% CI], large: 52.78 cm/s [50.62,54.91], small: 41.91 cm/s [39.67,44.11]; Wilcoxon signed-rank test: Z=707.00, p= 2.3×10^-19^). When analyzed separately for each of the 5 trial blocks, we observed that running speed was low in the first trial block and increased in the second trial block for both large and small reward conditions (figure 2a, right). Subsequently, running speed remained elevated for the large reward trials and decreased for the small reward trials. No difference between reward conditions was observed for the running speed from the reward platform back home (figure 2b, mean [99% CI], large: 43.70 cm/s [40.84,46.68], small: 43.33 cm/s [40.64,46.09]; Wilcoxon signed-rank test: Z=5056.00, p=0.74). Rats also spent more time consuming the reward on the platform at the end of the target arm that was associated with large reward as compared to small reward (figure 2c, mean [99% CI], large: 49.64 s [48.05,51.35], small: 6.41 s [5.88,6.97]).

**Figure 2:**
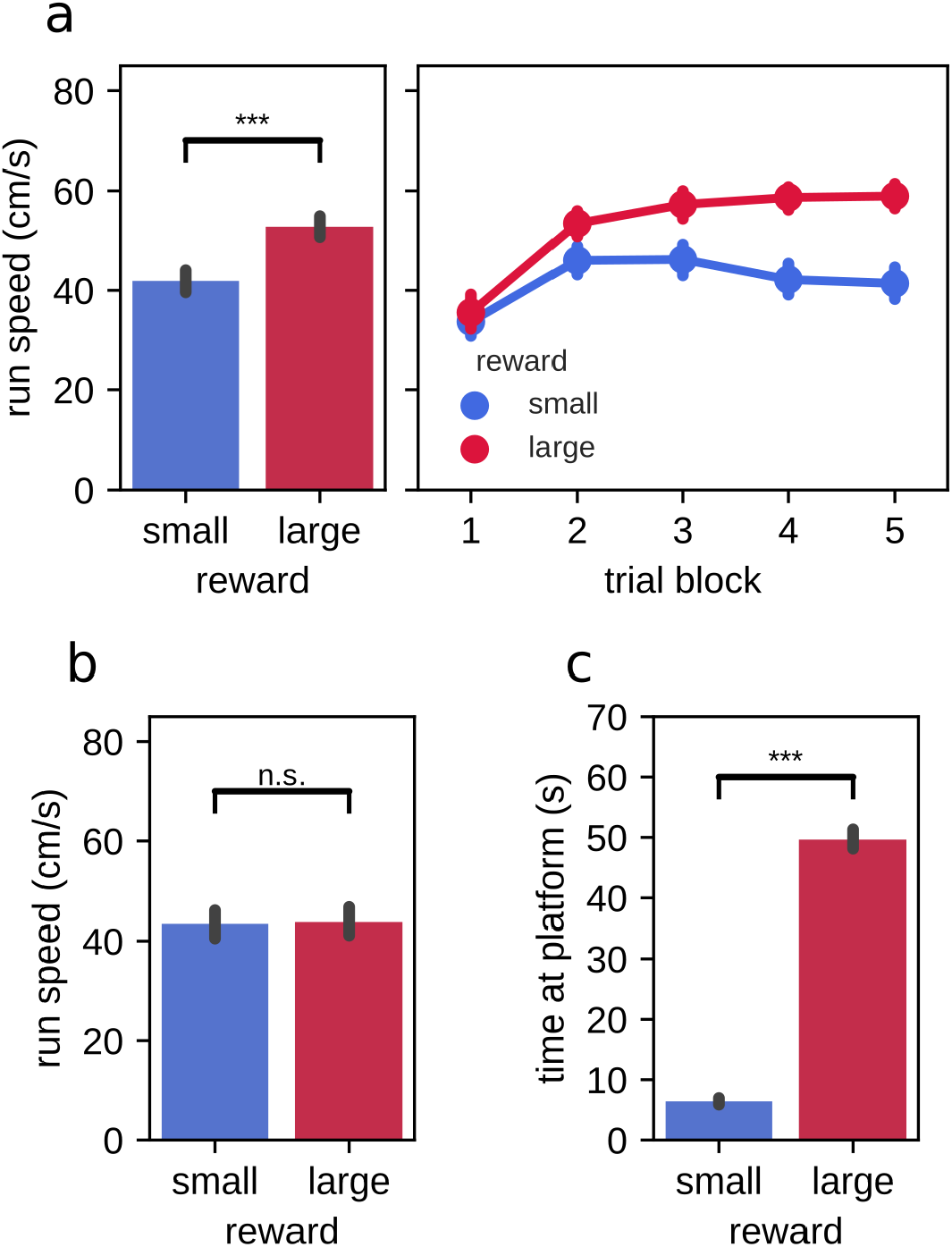
Run speed and time spent at the reward platform during instruction trials. (a) Run speed from home to the reward platform. Left: trial-averaged run speed for small and large reward targets. Right: per trial averaged run speed for small and large reward targets. (b) Trial-averaged run speed from the reward platform to home for small and large reward targets. (c) Trial-averaged time spent at the small and large reward platforms. Error bars represent the 95% confidence interval. ***: p<0.001.

### Stronger behavioral bias towards target arms associated with large reward amount after 2h delay

Overall, 21 rats perform a total of 151 sessions of probe test following 2h of delay. Rats made a median of 8 arm visits (inter-quartile range: 6-10, 151 sessions in 21 animals). There was a small tendency for rats to make more visits in the large reward environment (figure 3a; mean large-small difference [99% CI]: 0.53 [0.01,1.07]; Wilcoxon signed-rank test: Z=3065.50, p=0.015). Overall, we observed a preference for visits to the target arm over visits to the distractor arms in both large and small reward environments. On average, the probability to visit the target arm *p*(*T*) in both reward conditions (figure 3c; mean [99% CI], large: 0.47 [0.44,0.50], small: 0.35 [0.33,0.38]) was significantly better than chance under a binomial distribution *B*(*n*, *p*=0.25), where *n* was taken as the actual number of arm visits in each session (10000 simulations; Monte-Carlo p-value, large: p=0.0001, small: p=0.0001). On their first journey, rats were more likely than chance level (p=0.25) to visit the target arm (*p*(*visit*_1_=*T*); mean [99% CI], large: 0.81 [0.73,0.89], small: 0.62 [0.52,0.72]; binomial test under the null hypothesis of uniform arm visit probability, large: p= 7.1 × 10^-48^, small: p= 4.7 × 10^-22^) (figure 3b). The preference for the target on the first visit was stronger for the large reward environment than the small reward environment (mean large-small difference [99% CI]: 0.19 [0.07,0.32]; McNemar test, *H*_0_: *p_small_* = *p_large_*, *χ*^2^ =13.00, p=0.00011). Moreover, *p*(*T*) was significantly higher in the environment that was associated with large reward amount as compared to environment associated with small reward amount (figure 3c; mean large-small difference [99% CI]: 0.11 [0.08,0.15]; Wilcoxon signed-rank test: Z=1341.00, p= 4.7 × 10^-14^).

**Figure 3:**
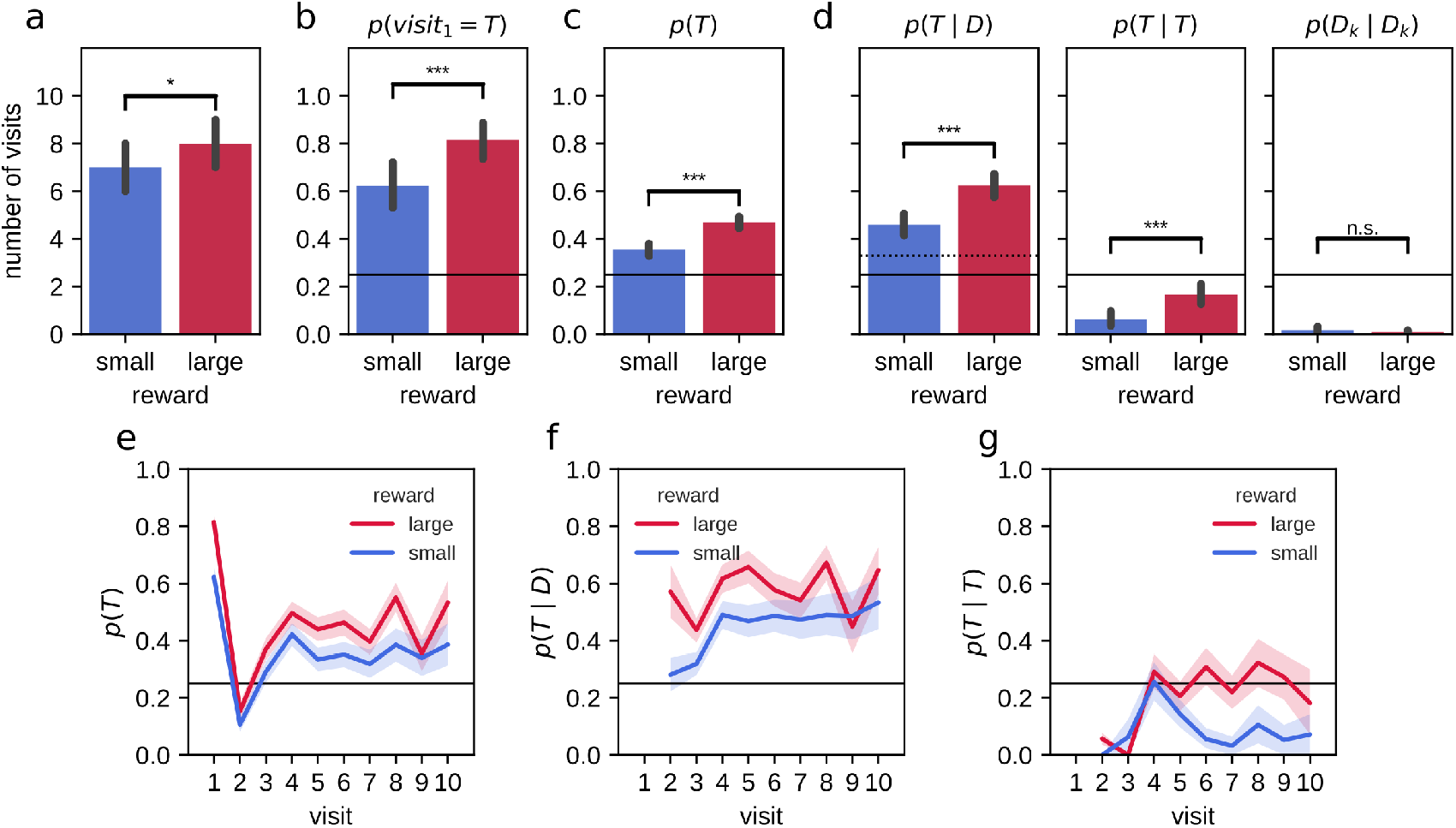
Stronger bias for large reward-place associations after 2h delay. (a) On average, animals made more arm visits in the 2-minute probe trial in the large reward environment as compared to the small reward environment. (b) The probability that animals first visited the target arm in the 2-minute probe trial was higher in the large reward environment as compared to the small reward environment. Performance for both large and small reward environments are well above chance. (c) The average probability of visiting the target arm in the 2-minute probe trial. (d) The associated conditional probabilities of visiting the target arm given that the rat previously visited a distractor arm (left) or the target arm (middle; i.e. repeat visit to the target arm) and the probability for a repeat visit to the same distractor arm (right). (e) Probability of visiting a target arm is higher across visits in the large reward environment compared to the small reward environment. On the first visit, animals often go straight to the target arm followed by a visit to a distractor arm on the second visit. On the remaining visits, the probability to visit the target arm remains constant. (f) The conditional probability of visiting the target arm given that the animal previously visited a distractor arm accross visit is higher for large reward than small reward. (g) The probability of a repeat visit to the target arm is low for initial visits but increases afterward. For the large reward condition, this increase persists while for the small reward condition the probability of making a repeat visit declines. Error bars represent the 95% confidence interval; solid line: 0.25 chance level, dashed line: 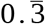 chance level, *: p<0.05; ***: p<0.001; n.s.: non-significant.

We looked in more detail at the behavior during the memory probe trial by separately analyzing the target arm preference for the first arm visit and subsequent visits. On the second visit, rats had a higher tendency to explore non-target arms, before a clear preference to revisit the target arm established in the remainder of the two minute probe test (figure 3e). We computed the conditional probability *p*(*visit_k_* = *T* ∨ *visit*_*k*-1_ = *D*) = *p*(*T* ∨ *D*) where *T* indicates the visit to a target arm and *D* to a distractor arm. In both large and small reward conditions, the revisit probability *p*(*T* ∨ *D*) is significantly higher than naive chance level (0.25) and the more conservative chance level of 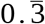 that assumes rats never immediately return to the exact same arm they just visited (figure 3d, left; mean [99% CI], large: 0.62 [0.58,0.67], small: 0.46 [0.41,0.51]). The target revisit probability is significantly higher for the large reward environment than the small reward environment (mean large-small difference [99% CI]: 0.17 [0.10,0.23]; Wilcoxon signed-rank test: Z=1913.50, p= 2.5 × 10^-9^).

Rats have a natural tendency to alternate maze arms and to avoid visiting the same arm twice in succession. Accordingly, rats made virtually no repeat visits to the same distractor arm (figure 3d, right; repeat probability *p*(*D_k_* ∨ *D_k_*); mean [99% CI], large: 0.01 [0.00,0.02], small: 0.02 [0.00,0.03]). However, rats did show an increased tendency to immediately return to the target arm without visiting any other arm in between, expressed as the conditional probability *p*(*T* ∨ *T*). While on average the probability *p*(*T* ∨ *T*) was lower than chance for both large and small reward conditions (figure 3d, middle; mean [99% CI], large: 0.17 [0.12,0.21], small: 0.06 [0.03,0.09]), they were significantly higher than the corresponding probability of repeat visits to distractor arms *p*(*D_k_* ∨ *D_k_*). Moreover, *p*(*T* ∨ *T*) was significantly higher for the large reward as compared to the small reward condition (figure 3d, middle; mean large-small difference [99% CI]: 0.11 [0.06,0.16]; Wilcoxon signed-rank test: Z=426.50, p= 5.9 × 10^-7^). Interestingly, an increase of the repeat visit probability *p*(*T* ∨ *T*) developed from the fourth visit in both small and large reward conditions, but the increase was only temporary for the small reward condition (figure 3g).

### Influence of spatial configuration

We next tested if other factors also contributed to the behavioral performance. Due to asymmetries in the configuration of target and distractor arms, the spatial configurations experienced varies between sessions. We quantified the behavioral performance in the memory probe trial separately for central, intermediate and edge target arm locations (see figure 4b. The general reward-seeking behavior in a memory probe trial did not vary systematically with the location of the target arm (figure 1a inset). Indeed, Poisson regression analysis confirmed that the location of the target arm did not significantly affect the total number of visits (*slope_small_*: −0.068 [-0.14,0.011], *slope_large_*: −0.02 [-0.091,0.066]).

**Figure 4:**
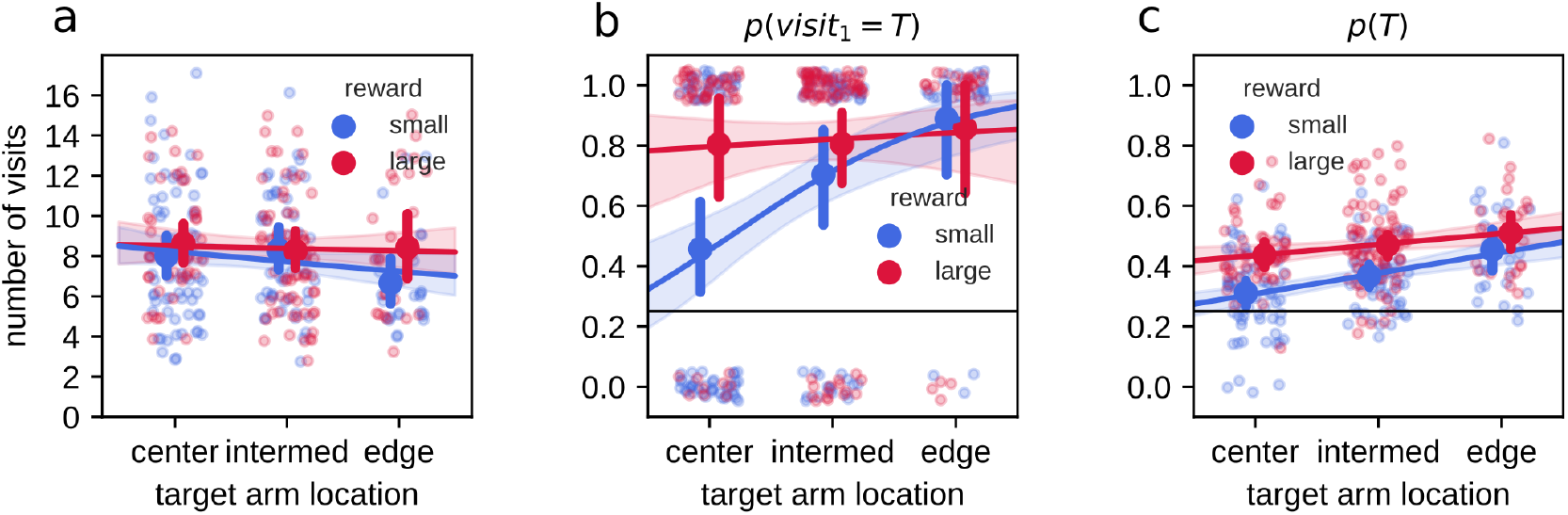
Effect of target arm location. In all panels, individual data points are shown as small semi-transparent dots (with added jitter to reduce overlap), filled circles represent the mean and 99% confidence interval, line and shaded region represent the mean and 95% credible region of posterior predictive samples from Bayesian GLM model fit. (a) Number of arm visits in 2-minute memory probe trial split by location of the target arm. (b) Probability of visiting the target arm on the first visit in the 2-minute memory probe trial. (c) Average probability of visiting the target arm in the 2minute memory probe trial.

The probability of the rats to first visit the target arm *p*(*visit*_1_=*T*) was robust to the centrality of the target arm when associated with large reward, but increased from central to edge target arm locations in the small reward condition (figure 4b). The result of logistic regression showed a significant positive relation between arm centrality and *p*(*visit*_1_=*T*) only for small reward (*slope_small_*: 1.1 [0.59,1.7], *slope_large_*: 0.16 [-0.45,0.78]), as well as significant difference in the regression slopes between large and small reward (*Δslope_large-small_*: −0.96 [-1.8,-0.12]). The difference between large and small reward conditions was largest for central arms, whereas for edge arm locations performance did not differ between reward conditions (figure 4b; post-hoc two proportion z-test for large-small reward differences at center, intermediate and edge arm locations, with Bonferroni corrected p-value for three hypotheses, center: z=-3.73, p=0.00058, intermediate: z=-1.35, p=0.54, edge: z=0.35, p=1).

We observed that the average probability of visiting the target arm *p*(*T*) also increased from central to edge locations, but now for both large and small reward conditions (figure 4c). Ordinary least squares regression confirmed positive slopes (*slope_small_*: 0.071 [0.042,0.096], *slope_large_*: 0.036 [0.0062,0.066]), and indicated that there was a small difference in the slopes for large and small reward (*Δslope_large-small_*: −0.029 [-0.071,0.0084]).

The presentation order of the left and right environments in the instruction trials and memory probe trials was randomized across sessions. While the total number of visits (mean last-first difference [99% CI]: 0.34 [-0.23,0.87]; Wilcoxon signed-rank test: Z=3285.50, p=0.059) (figure 5a) and the on-target probability of the first visit *p*(*visit*_1_=*T*) during the memory probe (mean last-first difference [99% CI]: 0.09 [-0.03,0.21]; McNemar test, *H*_0_: *p_first_* = *p_last_*, *χ*^2^ =21.00, p=0.1) (figure 5b) were not affected by the presentation order, the average target visit probability *p*(*T*) was marginally lower for the environment tested first (mean last-first difference [99% CI]: 0.03 [-0.01,0.07]; Wilcoxon signed-rank test: Z=3968.50, p=0.033) (figure 5c).

**Figure 5:**
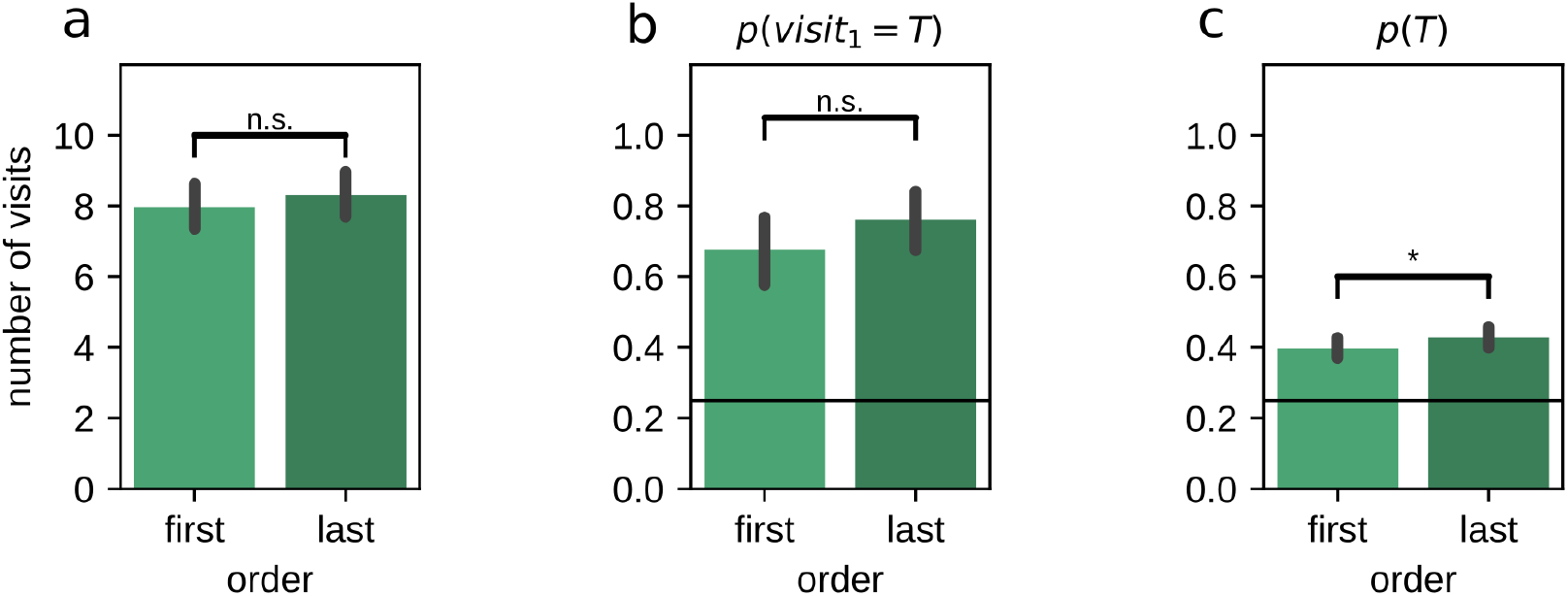
Effect of the order in which left/right environments were presented and tested. (a) Number of arm visits in 2-minute memory probe trial split by presentation order. (b) Probability of visiting the target arm on the first visit in the 2-minute memory probe trial split by presentation order. (c) Average probability of visiting the target arm in the 2-minute memory probe trial split by presentation order. Error bars represent the 95% confidence interval; *: p<0.05.

### Effect of long-term experience in the task

Rats were repeatedly trained in the same paradigm for several weeks, we thus asked if the animals’ performance varied over time. With increasing experience in the task, the total number of arm visits during the 2-minute memory probe trial decreased for both reward conditions (figure 6a, Poisson regression model, *slope_small_*: −0.019 [-0.027,-0.011], *slope_large_*: −0.022 [-0.03,-0.015]). This indicates that the animals reduced their reward-seeking behavior, possibly because they learned to recognize a memory probe trial that is never rewarded. The probability *p*(*visit*_1_ = *T*) increased significantly across sessions, but only for the small reward condition (figure 6b, logistic regression model, *slope_small_*: 0.075 [0.027,0.13], *slope_large_*: 0.037 [-0.014,0.1]). In contrast, the decrease in total number of visits did not affect the average probability *p*(*T*) to visit the target arm (figure 6c, ordinary least squares regression model *slope_small_*: −2.6 × 10^-5^ [-0.0025,0.0028], *slope_large_*: −5 × 10^-5^ [-0.0028,0.0026]).

**Figure 6:**
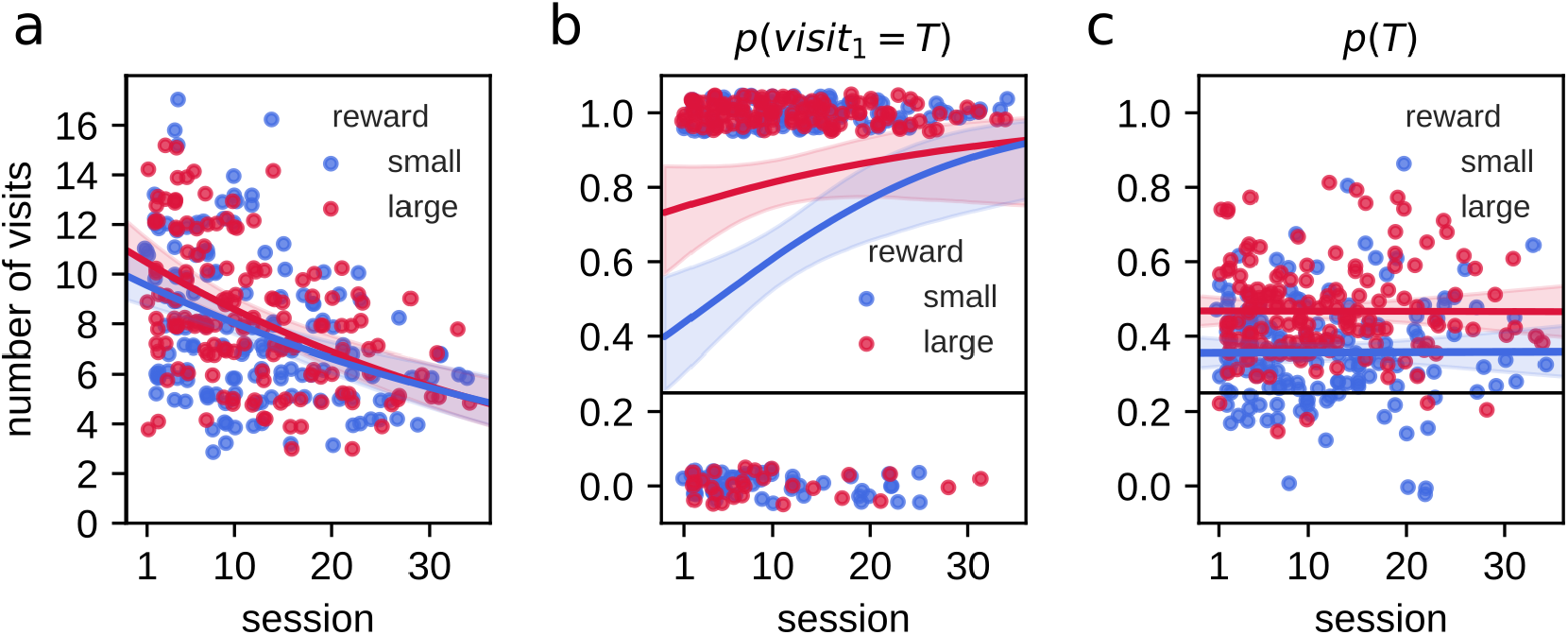
Effect of accumulated experience on behavioral performance. In all panels, individual data points are shown as small dots (with added jitter to reduce overlap), line and shaded region represent the mean and 95% credible region of posterior predictive samples from Bayesian GLM model fit. (a) The number of arm visits in 2-minute memory probe trial decreases as a function of experimental session for small and large reward environments. (b) The probability of visiting the target arm on the first visit in the 2-minute memory probe trial increases with experience only for the small reward condition. (c) The average probability of visiting the target arm *p*(*T*) in the 2-minute memory probe trial does not change with experience.

### Difference in performance between large and small reward conditions partially maintained after 20h delay

Rats were also tested for their memory of the target arms locations after 20h delay (9 animals, 59 sessions). They made similar number of total visits within the 2-minute probe test in large and small reward environments (figure 7a, mean large-small difference [99% CI]: −0.15 [-0.95,0.61]; Wilcoxon signed-rank test: Z=613.50, p=0.82). Overall, the animals displayed a preference for visiting the target arm associated to large reward amount compared with small (figure 7c; mean large-small difference [99% CI]: 0.06 [0.02,0.11]; Wilcoxon signed-rank test: Z=314.00, p=0.00023), with an average probability to visit the target arm *p*(*T*) higher than chance in the large reward condition but not the small reward condition (10000 simulations; Monte-Carlo p-value, large: p=0.0001, small: p=0.14). On their first journey, the probability of visiting the target arm *p*(*visit*_1_ = *T*) was higher than chance level (p=0.25; mean [99% CI], large: 0.53 [0.36,0.69], small: 0.41 [0.25,0.58]; binomial test under the null hypothesis of uniform arm visit probability, large: p=6.2 × 10^-6^, small: p=0.0097) Despite a tendency for the probability of first visiting the target arm associated to the large reward to be higher than for the small, the difference between the two reward conditions was not significant when including the probe memory test sessions only (mean large-small difference [99% CI]: 0.12 [-0.14,0.37]; McNemar test, *H*_0_: *p_small_*=*p_large_*, *χ*^2^ =14.00, p=0.31). However, the conditions in which the rats choose the first arm to visit are virtually the same between probe and routine test sessions. When combining all test sessions, the difference between reward conditions was then significant (probe test and routine, 116 sessions; mean large-small difference [99% CI]: 0.18 [0.01,0.35]; McNemar test, *H*_0_:*p_small_*=*p_large_*, *χ*^2^ =23.00, p=0.014)(figure 7b).

**Figure 7:**
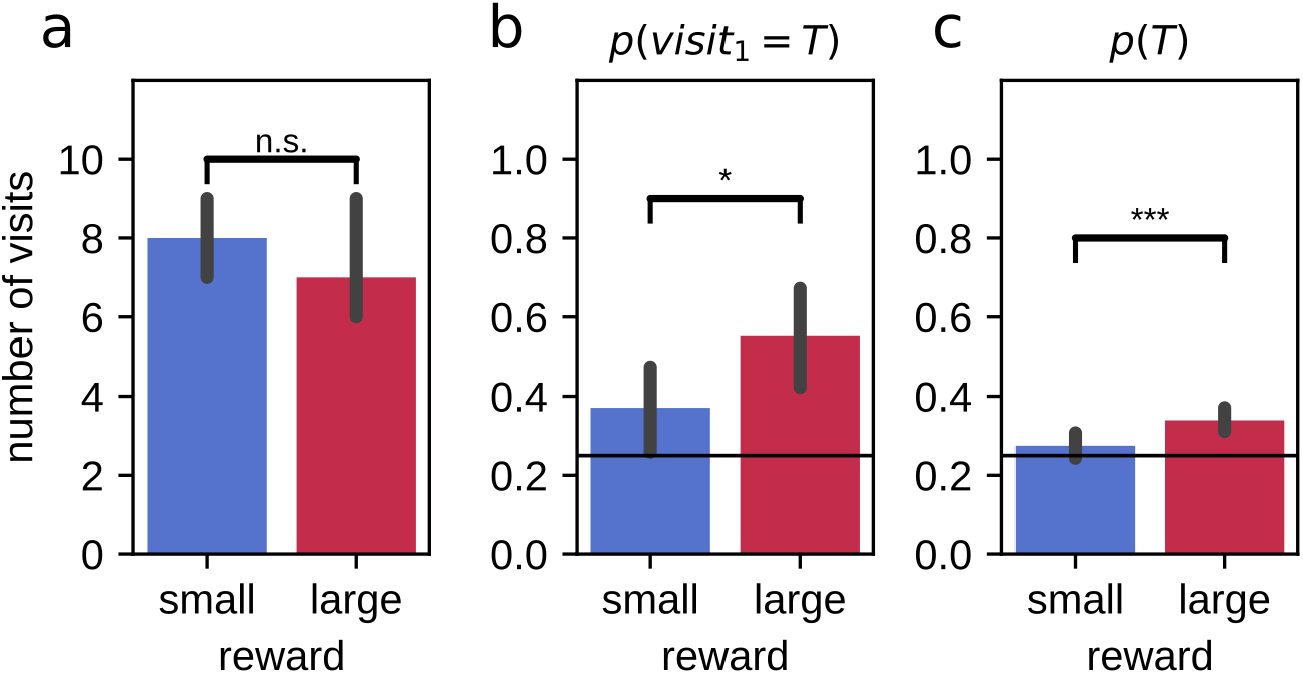
Difference in performances between large and small reward environments are partially maintained after a 20h delay. (a) On average, animals make more arm visits in the 2-minute probe trial in the large reward environment as compared to the small reward environment. (b) For all test sessions (probe test and routine) combined, the probability that animals first visit the target arm in the 2-minute probe trial is higher in the large reward environment as compared to the small reward environment. Performance for both large and small environments remains above chance. (c) The average probability of visiting the target arm in the 2-minute probe trial is higher in the large reward condition compared to the small reward condition. Performance for large reward, but not for small reward, is above chance. Error bars represent the 95% confidence interval; solid line: 0.25 chance level, dashed line: 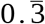 chance level, *: p<0.05; ***: p<0.001; n.s.: non-significant.

## Discussion

Our aim was to develop a behavioral paradigm to study the enhanced memory retention of salient experiences in rodents. Each day, rats were trained to learn a different food-place association on two contextually distinct semi-radial arm mazes. One arm was associated with a large reward amount, and the other a small amount of reward. The locations of the arm and the reward associated to the two environments were pseudo-randomly assigned every day. After a delay of 2h or 20h the animals were placed again in the two environments separately to be tested on their memory for the previously rewarded locations.

During training, rats showed a rapid increase in average run speed on journeys towards the reward. Moreover, a difference in speed developed throughout the training. The speed on journeys to the large reward remained stable while the speed to the small reward decreased towards the end of the training sessions, suggesting different motivational states consistent with the two different rewarded outcomes. These results indicate that the animals were able to quickly learn the food place associations.

The performance of the animals to retrieve the previously rewarded locations was assessed by monitoring the pattern of arm visits followed during the test phase after 2h and 20h delays. For both delays and both reward conditions, the rats were more likely than chance to first visit the target arm, which indicates that they remembered the food-place associations. Their probability of visiting the target arm throughout the 2min probe test period was also above chance level for all conditions except for the arm associated with small reward after 20h, possibly reflecting a lesser degree of confidence on their memory for the location of the rewarded arm in this condition. Consistent with the reported negative effect of prolonged delay on memory retention (Murre and Dros 2015), the animals’ performance was lower after a longer delay of 20h for both reward conditions. On average, the performance of the animals was also higher for the large reward location compared to small for both delays. These results are consistent with the studies carried in human showing a selective enhancement of memory for salient experiences, including the expectation of a higher value outcome, after both a short or long retention delay (Igloi et al. 2015; Gruber et al. 2016; Studte, Bridger, and Mecklinger 2017). Moreover, after 2h delay, we observed that overall the animals directly repeated visits to the target arm with a lower probability than expected by chance. However, they revisited more often for the target arm in the large reward condition as compared to the small reward condition. This suggest that a larger rewarding outcom e drove the animals away from their natural alternating behavior. Finally the results also validates the use of the dual reward-place associations task to study the mechanisms underlying the selective retention of memory in rodent.

Further analysis revealed that other aspects of the experience in the paradigm influenced the performances of the animals after 2h of retention delay. The performances of the animals varied in function of the location of the baited arm relative to the edge of the environments. The more radially distant the arm was from the edge the more the performances of the animals decreased. This effect was more pronounced in the small reward condition, so that performances for the arms close to the edges were similar between the two reward conditions and the difference progressively to be maximal for the arms most centrally located. First, these results confirm a modulation of memory retention by the amount of reward as the differences in performance cannot solely be explained by different seeking strategies related to motivational state. Second, it indicates that the reward-related enhancement of memory was interacting with other features of the experience potentially dependent on different memory systems (Ekstrom, Arnold, and Iaria 2014; Kirch et al. 2015).

Despite being highly familiarized with the majority of the task parameters, the performance of the animal improved over time, at the scale of weeks of training (meta learning). The improvement in performance may reflect learning associated with the changes introduced at the start of the experimental procedure, such as the increased retention delays, the different rewarded outcomes or the non-rewarded trials used as probe memory tests. The increase in memory performance was accompanied with a reduction of the seeking behavior during the probe tests, suggesting that the animals had adjusted their behavior to the fact that these trials were unrewarded, which argues in favor of the fact that at least part of the meta learning reflected the learning of the changes in the tasks. The meta learning related to memory performance was mainly observed for the small reward condition, while performance in the large reward conditions were already close to their maximum from the early phase in the training. This suggests that higher value outcome during training also accelerates meta learning.

The dual reward-place association task is suited to study the modulation of memory retention by features of experience as was demonstrated with varying the amount of reward associated to two similar but distinct experiences. However, the implementation of novel features would improve its quality, optimize the data collection and facilitate running the experiment. Currently, most aspects of the task, such as positioning the doors or reward delivery, are manually handled by the experimenter. The automation of these aspects would facilitate running the experiment as well as increase its reliability. The enhancement of memory for highly rewarded experience was dependent on locations presumably requiring the use of an allocentric strategy. Maximizing the number of locations requiring higher level of spatial integration, for example by increasing the total number of possible arm locations, would then optimize data collection. One confounding factor in the paradigm is the fact that the animals spent more time consuming the large amount of reward compared to small. Several approaches could be used to at least mitigate a putative effect of time spent at the reward location on memory retention: deliver different amount of reward at the starting point, common to both environments; use the natural behavior of rats to carry food to consume it in less exposed conditions or use different concentrations of rewarding agents in solution (Salvetti, Morris, and Wang 2014; Whishaw and Dringenberg 1991).

The choice of behavioral assays is critical, not only to ensure a behavioral read out of the cognitive process of interest, but also as it participates in optimizing the amount and reliability of the data collected. Experiments combining neuronal recording and/or manipulations with behavior are critical to understand how brain activity supports cognitive functions but they require heavy investments per animals. Repetitive and behaviorally constrain paradigm have the potential to increase the outcomes from experiments correlating brain activity with behavior by reducing variability and increasing the number of datasets collected per animal. Moreover, experiment involving manipulations, for example of particular aspects of an experience or directly of brain activity, benefits from the ability to compare the effects with internal controls from the same animals, between or within the same experimental sessions. The dual reward-place association paradigm is suitable for neuronal recordings and manipulations (Michon et al. 2019): the use of radial arms enables to render the behavior of the animals more stereotypical, the training can be repeated over weeks by every day changing the locations to be learned and it allows the use of internal controls between and within sessions.

## Author Contributions

FM and FK conceived and designed the study. FM, CK and JS carried the experimental work. FM, CK, JS and FK analyzed and interpreted the data. FM and FK wrote the manuscript. CK and JS reviewed the manuscript. FK coordinated and provided funding for the project.

## Acknowledgments

We thank Rudy D’Hooge for his comments on the design of the experiment.

